# Closed-loop Optogenetic Control in a Microplate Reader

**DOI:** 10.1101/2025.09.22.677797

**Authors:** Hari R. Namboothiri, Krishna Pochana, Bhavya Jaiswal, Azita Emami, Chelsea Y. Hu

## Abstract

Optogenetics integrates living cells and electronics into powerful cell–silicon systems, but prototyping their dynamics remains challenging. Current tools either require robotic liquid transfers into flow cytometers or rely on custom sensors with narrow dynamic range that limit controller performance. Additionally, current successful optogenetic feedback controllers only operate in chemostats or microfluidic devices that enforce constant growth, because models for growth-aware controller design in batch culture are lacking. Here we present LEMOS, a low-cost LED-embedded microplate that runs inside a commercial microplate reader. Coupled to a growth-aware multiscale model of gene expression for controller tuning, this platform enables rapid design-build-test-learn cycles for cell-silicon systems. We demonstrate closed loop setpoint tracking of gene expression in batch cultures within a standard microplate reader and show how growth dynamics complicates controller selection and tuning. Together, this platform reduces setup overhead and speed up iteration, enabling accurate real-time optogenetic feedback control.

## Introduction

Living cells, shaped by evolution, sense a wide diversity of molecules and convert these signals into coordinated gene expression and synthesis of complex organic compounds. Yet cellular responses are slow, noisy, and context dependent, which makes precise regulation difficult. Electronic systems have limited direct access to chemicals but excel at high-speed signal transduction, computation, communication, and control. Combining the two yields cell-silicon hybrids that pair biochemical breadth with engineered precision, enabling remote sensing and adaptive biomanufacturing. Development of such an integrated cell-silicon system has become possible through the establishment of bidirectional communication: light can modulate gene expression through optogenetic control, while fluorescent or luminescent protein reporters can relay gene expression readouts as electronic signals through photodiodes(*1-3*).

Optogenetics uses light to control cellular processes, enabling rapid, reversible, and precise spatiotemporal regulation of gene expression, capabilities that chemical inducers often lack(*4*). Such control has been implemented at the transcriptional(*5-9*), translational(*10, 11*), and post-translational levels(*12-14*). For example, the classic light-responsive CcaSR system comprising a photoreceptor histidine kinase and a response regulator enable reversible activation and repression of transcription(*6*). A remaining practical hurdle for cell-silicon control in typical labs is coupling real-time monitoring with dynamic illumination in standard microplate readers. Prior solutions achieve closed-loop regulation using chemostats with robotic sampling(*15, 16*) or single-cell feedback via automated microscopy in a mother-machine(*17-19*). However, the complexity and cost of these platforms limit adoption and slows the design-build-test-learn cycle. To overcome these issues, Steel, H. et al.(*20*) developed a custom miniaturized chemostat that enabled closed-loop optogenetic control, though the device’s utility is curbed by the sensor’s limited dynamic range. Additionally, these platforms maintain steady-state growth, which eliminate the growth dynamics that arise in batch and other non-constant growth conditions, constraining applicability.

The most common assay in synthetic biology grows cultures in well plates and uses a commercial microplate reader to measure optical density (OD) and fluorescence. External LED-array stimulators have been built to modulate light(*21-25*), but typically require a human operator(*26*) or a robotic stage to transfer to a microplate reader(*27*) for measurement. Several all-in-one platforms combine illumination with onboard fluorescence and OD_600_ sensing for closed-loop control, yet performance is limited by the sensing hardware(*28, 29*). RT-OGENE uses a digital camera on a stationary stage to read fluorescence and OD_600_, which narrows dynamic range. The system also does not allow shaking, reducing growth and sustained expression(*28*). The optoPlateReader moves forward with an LED and photodiode sensor and allows shaking, but it supports only a single reporter and has limited sensitivity for both fluorescence and OD_600(*29*)_. Both studies attempted feedback control; however, sensor limitations and the lack of an accurate gene expression model constrained controller performance. As a result, progress toward practical cell-silicon hybrid systems has been limited, and there remains a need for an easy-to-use, reader-compatible platform for closed-loop studies in batch culture that accelerates the DBTL cycle.

In this work, we built the LED-Embedded Microplate for Optogenetic Studies (LEMOS) device as a prototyping platform and demonstrated its utility in the DBTL cycle for cell-silicon hybrid dynamical systems. LEMOS is a low-cost (≈$140), LED-embedded microplate that operates inside a commercial microplate reader and synchronizes illumination with measurement by blanking LEDs during reads. In batch *E. coli* cultures, we first mapped open-loop responses, then showed that proportional control tracks predefined setpoints, although with notable overshoot. Using our growth-dependent gene expression framework (GEAGS)(*30*), we found that the observed dynamics arise from growth-dependent dead time in gene expression and predicted that adding integral and derivative actions would improve performance. Guided by the model, we designed and implemented Proportional-Integral (PI) and Proportional-Integral-Derivative (PID) controllers that improved both accuracy and response speed across conditions. By enabling feedback without leaving the microplate reader, LEMOS makes *in silico* feedback control practical in batch culture, lowers the barrier to rapid DBTL, and is readily extendable to other bacterial strains and optogenetic mechanisms.

## Results

### LEMOS Device Design and Validation

The LEMOS device is designed to match the layout of a standard 96 well microplate, ensuring compatibility with conventional microplate readers (Fig. 1A). The 3D printed frame houses 16 slots for polydimethylsiloxane (PDMS) microwells, each with an individually addressable light-emitting diode (LED) mounted on the inner wall. The LEDs are controlled by an Arduino Nano33 IoT microcontroller with built in Bluetooth, enabling wireless communication with an external computer. The microcontroller and the LEDs are powered by a Li-ion battery housed within the device, connected to the charging module and a three-position switch that toggles between ‘On’, ‘Off’ or ‘Charging’ modes. The detailed electronic components can be found in Supplementary Information (SI) Section 1 and the operation of the electronic components is described in Methods. To ensure sterility during cell culture experiments, PDMS sleeves are cast for each use via soft lithography (see Methods). PDMS is a transparent, biocompatible elastomer well-suited for microwell fabrication due to its flexibility, chemical inertness, and ease of molding. Its optical transparency also enables fluorescence measurements during experiments. A 3D-printed mold (Fig. 1C) was inserted into the LEMOS frame during casting (Fig. 1B) to form wells shaped for incubation of 200 µl bacterial cultures (Fig. 1D).

**Fig. 1.**
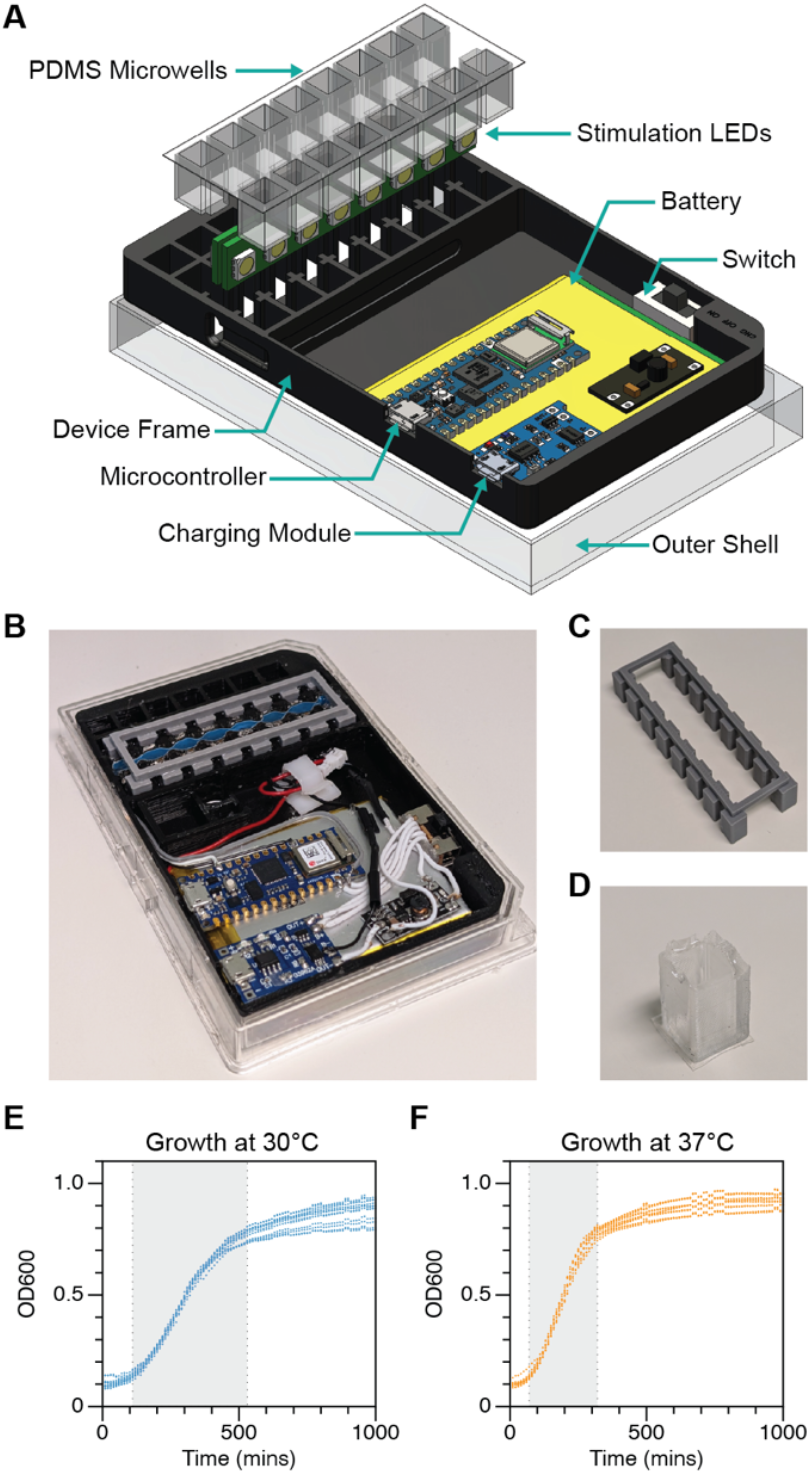
LEMOS device construction and cell growth characterization. (A) Exploded schematic of the LEMOS device showing PDMS microwells, stimulation LEDs, battery, switch, microcontroller, charging module, device frame, and outer shell. (B) Assembled device with the 3D-printed mold used to cast the PDMS microwells. (C) 3D-printed mold for microwell casting. (D) PDMS microwell casted in the LEMOS device (E) Growth curves of *E. coli* in LEMOS at 30 °C and (F) 37 °C (N=12 technical replicates). Gray shading marks the logarithmic (exponential) growth phase.

To assess whether LED-generated heat affects cell growth, we monitored bacterial cultures over 16 hours in the LEMOS device using continuous OD_600_ measurements in a standard microplate reader at both 30 °C and 37 °C. As shown in Figs. 1E and 1F, cell growth at 37 °C indicating that heat emitted by the LEDs had no apparent impact on its growth condition. Cultures at 30 °C exhibited a prolonged exponential phase compared to those at 37 °C (Figs. 1E and 1F). Therefore, all subsequent experiments in LEMOS were conducted at 30 °C to maximize the dynamic measurement window before stationary phase onset.

### System Characterization and Open-Loop Optogenetic Control

We used the CcaSR two-component system (TCS) optogenetic regulator system to control expression of superfolder green fluorescent protein (sfGFP). The system consists of CcaS, a membrane-bound histidine kinase that senses green light, and CcaR, the corresponding response regulator(*6*) (Fig. 2A). Upon green light exposure, CcaS autophosphorylates and transfers the phosphate to CcaR, which dimerizes to activate the PcpcG2 promoter and induce sfGFP transcription. Red light reverses this process by promoting CcaS dephosphorylation, thereby repressing expression. To characterize the system, we performed a bulk fluorescence assay in a 96-well microplate. Cultures were incubated at 37 °C under specific light conditions in a custom light box, and fluorescence was measured manually every hour, for six hours, using a microplate reader. The system functioned as expected in *E. coli* MG1655, with sfGFP activation under green light, repression under red, and intermediate expression in the dark due to basal promoter activity (Fig. 2B).

**Fig. 2.**
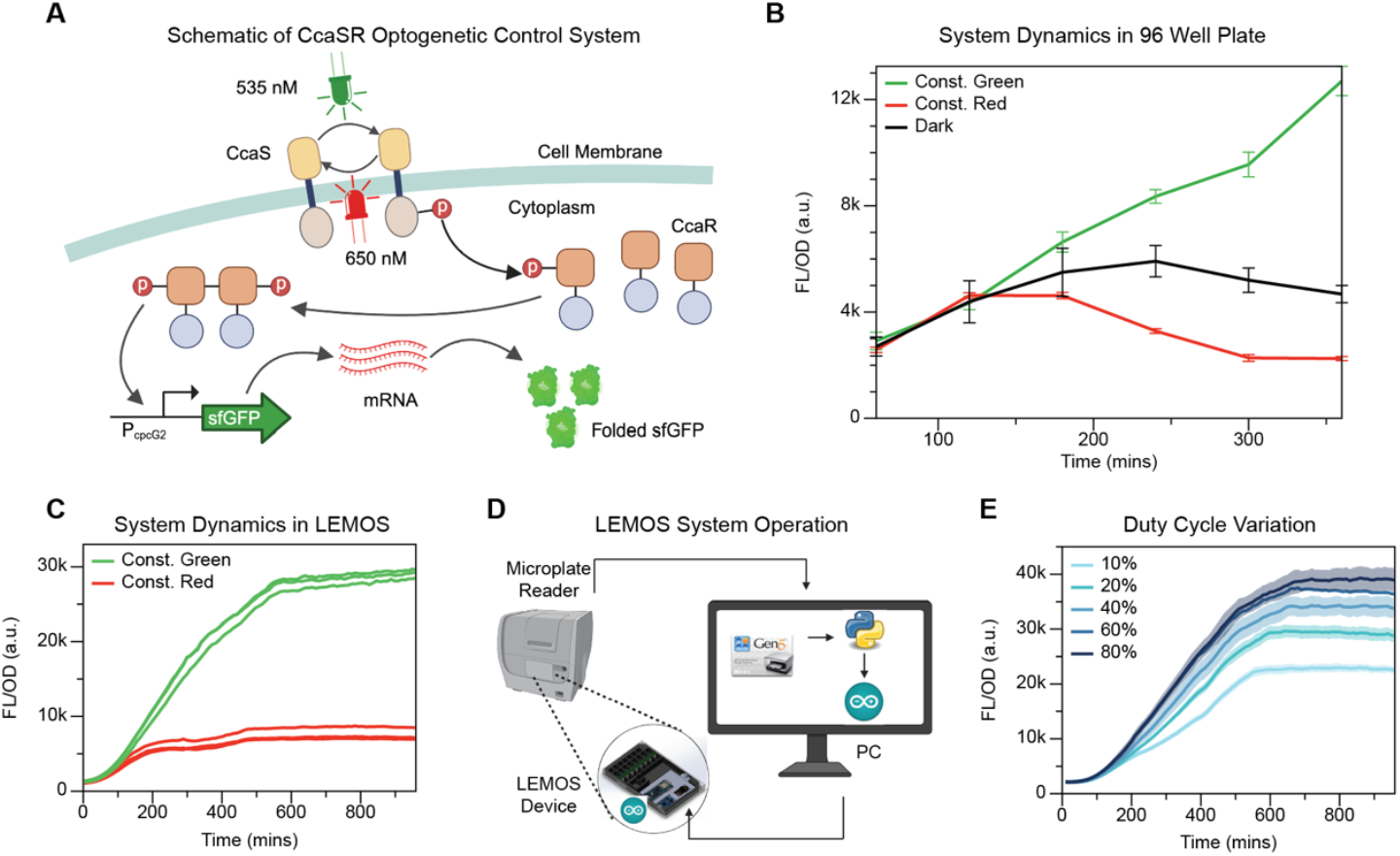
Optogenetic system characterization in LEMOS and open-loop control dynamics. (A) CcaSR two-component optogenetic system: green light (535 nm) activates CcaS/CcaR to upregulate PcpcG2-sfGFP expression, and red light (650 nm) downregulates the expression. (B) Activation-repression dynamics in sfGFP fluorescent/OD_600_ in a standard 96-well plate (n=4) under LED illumination; the plate is manually transferred to the microplate reader for measurement six times over six hours. (C) Same assay in the LEMOS device (N=3) for 16 hours, with one measurement every 10 minutes. (D) LEMOS operation schematic. During the 16-hour time course, the device remains in the microplate reader. The reader measures GFP fluorescence and OD_600_ and streams data to a computer, which handles timekeeping and operates Arduino routines that command the on-board microcontroller to temporarily disable LED illumination during each measurement. (E) Open-loop responses in LEMOS to varying green-light duty cycles (10-80%, N=3). Solid lines indicate means; shaded bands indicate standard deviations. N denotes the number of technical replicates; n denotes for the number of biological replicates.

To enable continuous measurement, we inoculated the same cell lines within the microwells of the LEMOS device and incubated it in a microplate reader. One key requirement for this setup was to prevent interference from the LEMOS LEDs with the optical measurements taken by the microplate reader. To address this, we ensured synchronization between the LEMOS device and an external computer, which handled timekeeping and communicated with the device via Bluetooth. The microplate reader was programmed to record OD_600_ and fluorescence every 10 minutes. During each interval, the LEDs were programmed to turn off during the first and last minute to avoid signal interference during measurements. As shown in Figs. 2C and 2D this setup allowed real-time monitoring of sfGFP expression dynamics under both green and red light over a 16-hour time course, with measurements taken every 10 minutes.

To evaluate the system’s response to actuating parameters, we characterized its dynamics under varying duty cycles. Duty cycle refers to the fraction of time within an interval during which green light is applied, with red light used for the remainder. Our control period is set to be 10 minutes, then the LEDs were on for 8 minutes every interval, during which we varied the proportion of green and red light. A 100% duty cycle corresponds to constant green light, while 0% corresponds to constant red light. As shown in Fig. 2E, we tested duty cycles ranging from 10% to 80%. The resulting sfGFP expression levels exhibited a clear positive correlation with duty cycle, indicating sensitive responses of system output (sfGFP signal) to the actuating parameter (duty cycle).

### Closed-Loop proportional feedback Control for Setpoint Tracking

Next, we implemented closed-loop cell-silicon feedback control for setpoint tracking using the LEMOS device. Setpoint tracking is a standard closed-loop task in chemical engineering that drives a measured system variable to a defined value. In this setup, as shown in Fig. 3A, gene expression in living bacterial cells serves as the process plant, with sfGFP fluorescence as the output. The microplate reader senses this output and transmits the data to an external computer acting as the controller. Based on the measured signal, the computer calculates control actions and sends commands to the LEMOS microcontroller, which adjusts the LED duty cycle (the actuator) to regulate gene expression via optogenetic input. In our case, the setpoint is a predefined FL/OD_600_ intensity that signifies the amount of sfGFP expressed per cell on average.

**Fig. 3.**
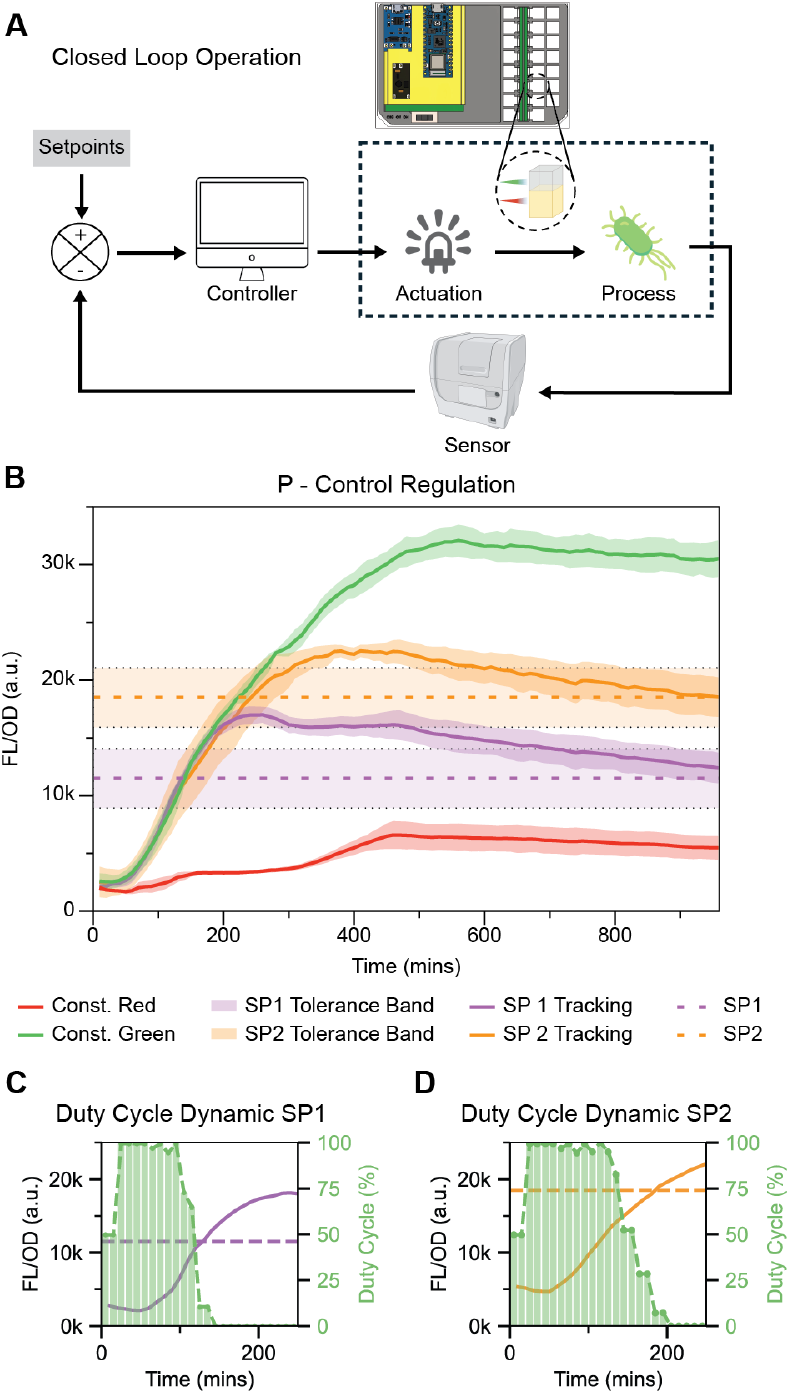
Closed-loop control in LEMOS. (A) Schematic of closed-loop operation: the microplate reader functions as the sensor that measures FL/OD, the computer functions as the controller, and the LEMOS device actuates LEDs to regulate gene expression. The process plant is gene expression in living bacterial cells. (B) Proportional (P) control tracking of fluorescence setpoints. Constant-light references are green and red (N=4, n=2). Setpoint runs over 2 independent days (N=9, n=2, total 18 trajectories). Solid lines indicate the mean FL/OD; shaded bands indicate standard deviations. Dotted lines mark setpoints (SP1 = 1.15×10^5^ a.u., SP2 = 1.85×10^5^ a.u.); shading around setpoints indicate ±10% tolerance. (C– D) Duty-cycle command for representative SP1 and SP2 runs; each panel shows a single well trajectory. N denotes the number of technical replicates; n denotes for the number of biological replicates.

To evaluate setpoint tracking, we assigned two distinct setpoints to two groups of microwells, corresponding to the orange and purple dotted lines in Fig. 3B. The controller continuously compared the measured fluorescence to each well’s assigned setpoint and computed the LED duty cycle for the next 10-minute control interval. We first implemented Proportional control (P-control), where the duration of green light exposure in each cycle (*t*_*green*_) is computed as:

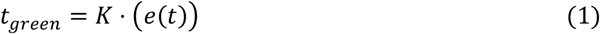

Here *e*(*t*) is the error between the measured FL/OD_600_ signal and the setpoint at time *t*, and *K* is the proportional gain that determines the controller’s response sensitivity. The remaining portion of the 8-minute control interval not occupied by green light was filled with red light. Because no measurement is available at *t* = 0, the controller initializes with a default 50% duty cycle for the first two intervals. Figs. 3C and 3D illustrated how the duty cycles change over time for the two setpoints. In these plots, the height of each green bar represents the fraction of green lights (i.e., the duty cycle) applied during each 8-minute control period. As expected, the duty cycle decreases rapidly as the measured sfGFP approaches the setpoint eventually reaching 0% once signal reaches the setpoint. As shown in Fig. 3B, both setpoints, FL/OD = 11.5e4 (SP1) and 18.5e4 units (SP2), were successfully tracked by the P-controller. However, despite the fast response time of the electronic actuators, both trajectories exhibited notable overshoot before gradually converging to their respective setpoints.

### Growth-Dependent Dead Time Leads to Signal Overshoot

To improve controller performance, we used a model-guided strategy to address a key question: does the consistent overshoot under P control arise from standard gene-expression kinetics or from an unexpected mechanism? To test this, we constructed a deterministic kinetic model of the system’s dynamics.

As shown in Fig. S7, a traditional growth-independent effective model fails to recapitulate the observed dynamics. To capture batch-culture behavior, we adopted the gene expression across growth stages (GEAGS) framework(*30*): we fit population growth with a logistic function (Fig. 4A) and modeled gene-expression rates as functions of the instantaneous growth rate. Model details are provided in SI section 4. With these modifications, the optogenetic GEAGS simulations closely matched the experiments, accurately reproducing sfGFP dynamics under both constant green and red-light inputs (Fig. 4B).

**Fig. 4.**
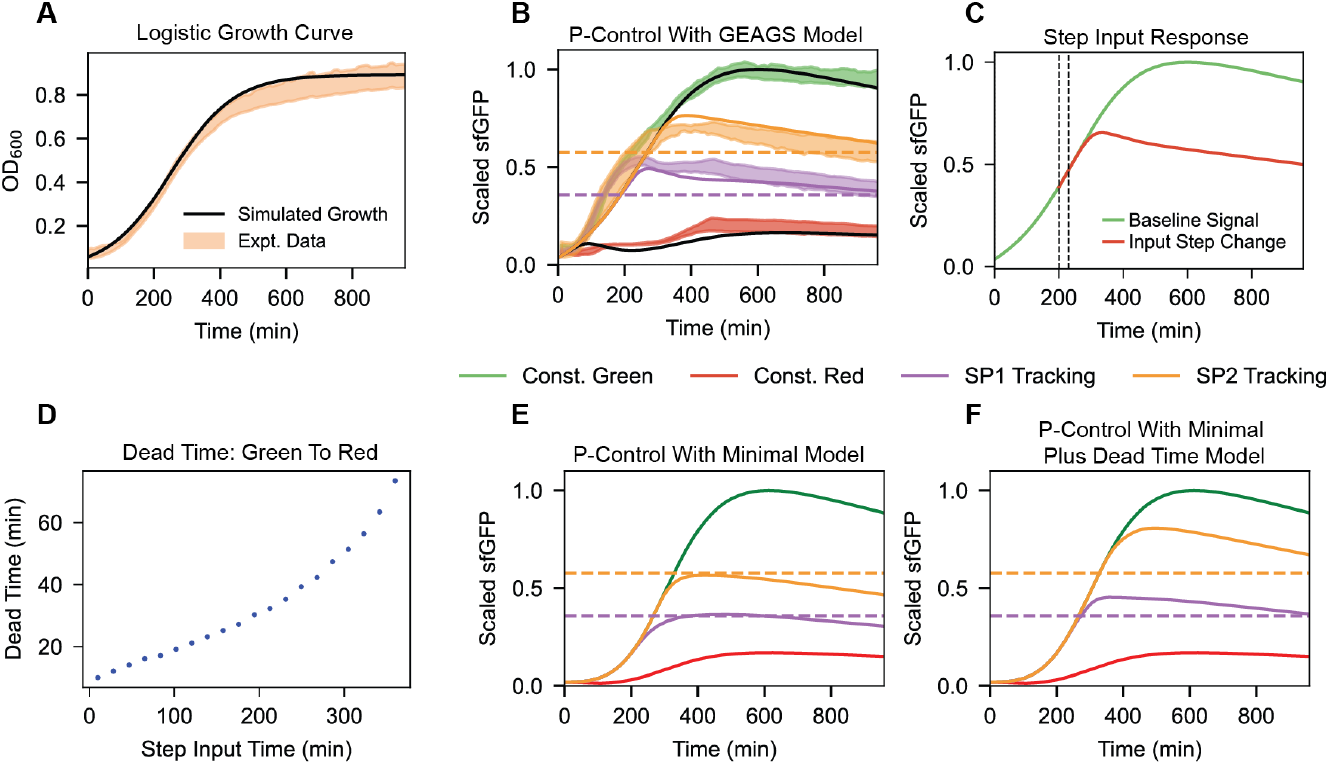
GEAGS growth-aware modeling explains dead time and P-control behavior. (A) Simulated logistic growth overlaid with measured OD_600_ (N=18, n=2). (B) P-control simulations with the GEAGS model overlaid on experimental setpoint-tracking data. Solid lines indicate model simulation, shaded regions indicate experimental data (N=9, n=2 for set point tracking). Dashed colored lines represent the predetermined setpoint. (C) Step-response simulation with GEAGS model showing dead time when switching the actuator light from green to red at t=200 min (baseline vs. input step change). (D) Response dead time of green-to-red switch increases over time. (E) P-control dynamics simulated with a minimal model omits gene expression dynamics associated time delay (SI section 5), showing minimal system overshoot. (F) P-control dynamics simulated with the minimal-plus-deadtime model, recapitulates the observed overshoot. N denotes the number of technical replicates; n denotes for the number of biological replicates.

To investigate the consistent overshoot observed in the P-control regulation experiment, we simulated the system using the GEAGS model framework(*30*). The simulation results closely matched the experimental data (Fig. 4B), capturing the overshoot behavior. This agreement suggests that the overshoot arises from well-understood mechanisms intrinsic to the system. We hypothesize that it is primarily driven by inherent delays in the gene expression process, such as translational and protein folding lag.

To quantify system delays, we measured dead time in the optogenetically controlled circuit.

Dead time is the interval after an input change during which the output shows no detectable response; here, the lag between an LED color shift and the resulting measurable change in FL/OD. We first simulated a baseline FL/OD trajectory under constant green light (Fig. 4C, green curve). We then introduced step changes from green to red at multiple time points and recorded how long it took the FL/OD signal to deviate from the baseline. At each time *t*, dead time was defined as the duration required to reach a 0.1% change from the baseline FL/OD (Fig. 4C). As shown in Fig. 4D, dead time increases over time, indicating a gradual decline in system responsiveness as the batch culture approaches the stationary phase. This trend is consistent with the general understanding that gene expression slows down as cellular resources are depleted during growth arrest, resulting in increasingly sluggish system dynamics.

To isolate the effect of dead time on closed-loop control dynamics we first simulated a delay-free minimal model that assumes minimal gene-expression delay and omits protein folding and two-component switching (equations S30-S32, SI Section 5). As expected, the closed-loop response showed negligible overshoot (Fig. 4E). To examine the role of deadtime in isolation, we then introduced a minimal-plus-dead time model by inserting the experimentally estimated dead time *t*_*dt*_(*t*) at each time step (Fig. 4D). Under P control, the controller at time *t*_*i*_ computes the duty cycle based on the sfGFP value at a prior time *t*_*i*_ ™ t_*dt*_(*t*_*i*_), emulating the delay and effectively shifting the perceived system state. This model reproduces the experimental overshoot (Fig. 4F), indicating that the overshoot arises from biological sluggishness: while sensing, computation, and actuation are millisecond-scale electronically, transcription, translation, and fluorescent-protein maturation limit the overall response.

### Implementation of PI and PID Feedback Control

Since proportional control consistently produced overshoot driven by dead time and slow gene-expression dynamics, we next evaluated proportional–integral (PI) and proportional– integral–derivative (PID) controllers. In control theory, PI control eliminates steady-state error by integrating past errors, while PID adds derivative action to improve transient response by anticipating changes in the error(*31, 32*). These strategies are particularly useful for slow or delay-prone systems, where proportional control alone is insufficient. We implemented both PI and PID schemes to test whether they could achieve more accurate setpoint tracking and faster convergence.

We first examined a PI controller to compensate accumulated error. The integral term sums past error (Eq. 2), which can reduce overshoot and improve tracking accuracy(*32*). In our implementation, the controller updated the duty cycle once every 10 minutes according to:

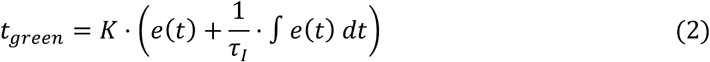

Here *τ*_*I*_ is the integral time constant, which scales the accumulated error e(t). Using the GEAGS model, we optimized PI controller gains and selected *τ*_*I*_ = 1500 min and *K* = 0.013 min/nM, Simulations showed that the PI controller markedly reduced overshoot and achieved convergence at both setpoints (Fig. 5A), in contrast to P control (Fig. 3B). We then implemented this model-guided design on the LEMOS plate, which confirmed consistently low overshoot for both setpoints (Fig. 5B). Tracking at SP2 was slightly less optimal, with minor overshoot. As indicated in Fig. 5E, the duty-cycle update appears insufficiently responsive, contributing to this residual overshoot and suggesting room for further controller refinement.

**Fig. 5.**
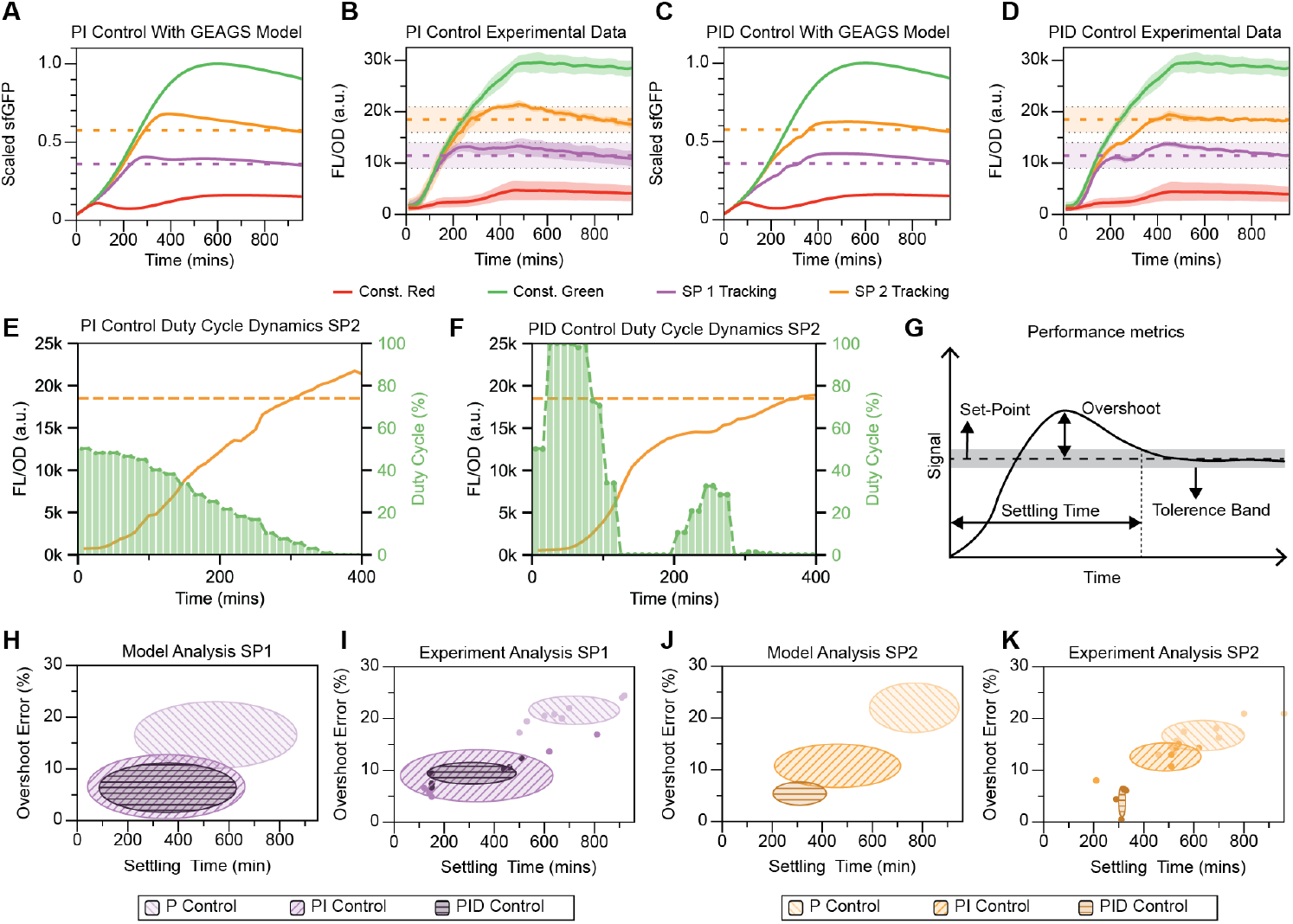
Model-guided PI and PID regulation. (A) GEAGS-based PI control simulations with tuned gains. (B) PI control experiments in LEMOS: constant-light references (N=14, n=6 each) and setpoint tracking at SP1 and SP2 (N=6, n=2 each). (C) GEAGS-based PID control simulations with tuned gains. (D) PID control experiments in LEMOS: constant-light references (N=14, n=6 each) and setpoint tracking at SP1 and SP2 (N=6, n=2 each). (E–F) Duty-cycle command for representative SP2 tracking with PI and PID controller; each panel shows a single well trajectory. (G) Definition of performance metrics: overshoot (%) and settling time to enter and remain within a ±10% band around the setpoint. (H–K) Comparison of P, PI, and PID performance from model predictions (H, J) and experiments (I, K) for SP1 and SP2. N denotes the number of technical replicates; n denotes for the number of biological replicates.

To further improve setpoint tracking performance, we implemented a proportional-integral-derivative (PID) control strategy. In addition to the proportional and integral terms, PID control includes a derivative term that responds to the rate of change of the error:

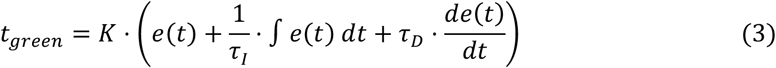

Here *τ*_*D*_ is the derivative time constant, which sets how much the slope of the error contributes to the control signal. The derivative term anticipates future error from its current rate of change(*32*). Using the GEAGS model, we optimized the PID controller parameters and selected *τ*_*I*_ = 1500 min, *K* = 0.032 min/nM, and *τ*_*D*_ = 120 min. Simulations showed that PID control (Fig. 5C) substantially reduced overshoot and improved convergence for SP2 relative to PI control, with only minor improvement for SP1. We implemented this design on the LEMOS device, and the experiments matched the predictions: PID enhanced tracking for SP2 with limited added benefit for SP1 (Fig. 5D). As shown in Fig. 5F, the duty-cycle trajectory exhibits a stair-step profile. An early rapid rise in signal (before ∼100 min) prompted a strong reduction in green light. As the signal’s rate of increase slowed, the controller raised the duty cycle between 200 and 270 min, followed by another sharp reduction near 300 min. This pattern is consistent with derivative action operating on a slow, sampled system, yielding anticipatory but discretely timed adjustments(*31*).

To compare controller performance, we defined two metrics (Fig. 5G). With activation at *t* = 0 and a 10% tolerance band around each setpoint, overshoot error is the maximum deviation of the sfGFP signal from the setpoint, and settling time is the time required for the signal to re-enter the tolerance band after the overshoot. We visualized performance on a map with overshoot on the y-axis and settling time on the x-axis, where strategies closest to the origin achieve faster responses and more accurate tracking.

For SP1, we evaluated P, PI, and PID control. In GEAGS simulations, we generated 50 trajectories by perturbing parameters ±15% around their fitted values and recorded the resulting metrics. The transition from P to PI control clearly improved both speed and accuracy, whereas PID yielded little additional benefit (Fig. 5H). The experimental analysis showed the same pattern (Fig. 5I). Applying the same procedure to SP2 in both simulations (Fig. 5J) and experiments (Fig. 5K) showed a consistent progression in performance from P to PI to PID.

Overall, these results indicate that derivative action was dispensable for SP1 tracking but became essential for SP2. We attribute this difference to growth-phase effects: SP1 was reached earlier, during logarithmic growth when system dead time was small and responsiveness was high, whereas SP2 was reached later, when the system was more sluggish, making derivative control critical for maintaining responsiveness.

## Discussion

Our work effectively showcases that LEMOS device makes closed-loop optogenetic control as routine as reading OD and fluorescence in a commercial microplate reader. By embedding LEDs in a microplate and blanking illumination during measurement windows, this platform preserves reader sensitivity and removes transfers between stimulation and readout. In batch *E. coli* cultures, proportional control tracked early setpoints but overshot as growth slowed. A growth-aware GEAGS model explained this behavior as growth-dependent dead time, and guided PI and PID designs that improved accuracy and settling across conditions.

Mechanistically, gene expression in batch culture behaves as a process plant whose effective delay increases as cultures approach stationary phase. Transcriptional and translational latencies, protein maturation, and biomass dilution slow the apparent response relative to the controller’s sampling period and setpoint changes. Since responsiveness depends on growth phase, a controller tuned for one setpoint or phase may be suboptimal in another. This highlights the multiscale nature of closed-loop gene regulation, where population dynamics shape molecular kinetics and determine which control structures and gains provide robust performance.

The LEMOS device makes closed-loop optogenetic control practical inside the same microplate reader used for routine OD and fluorescence, lowering the barrier to model-informed DBTL and revealing growth-phase effects that are often hidden in chemostats or single-cell rigs. Open-loop kinetics collected with LEMOS guided development of a growth-dependent deterministic model that accurately captures the hybrid system’s dynamics and guides controller design. To our best knowledge, this is the first end-to-end DBTL cycle to achieve functional closed-loop optogenetic control in batch culture.

Together, the accessible hardware and growth-aware model make DBTL for closed-loop cell-silicon systems a routine workflow, enabling rapid iteration and more reliable design. Both the device and the modeling framework are generalizable and can be extended to other hosts and optogenetic actuators.

The current LEMOS device offers three illumination channels (red, green, blue; RGB). In this work we used the green and red channels to drive the CcaSR system. The device is ready to evaluate systems using blue-light actuators, such as EL222(*33, 34*), iLID(*35*), or VVD(*36*). Variants can also be built to incorporate different LED lights to test optogenetic systems that are stimulated by, for example, near infrared light(*37*) and violet light(*38*). In addition, we will also use the platform to study layered control in synthetic biology(*39*) by combining molecular feedback or feedforward circuits with cell-silicon control, and quantify how layering affects setpoint tracking and disturbance rejection performance. On the control-algorithm side, because response speed varies with growth, controller gains should adapt accordingly. We will explore gain scheduling based on OD or online growth-rate estimates, add anti-windup and explicit disturbance-rejection tests, and, leveraging the calibrated GEAGS model(*30*), implement model predictive control (MPC) that uses short-horizon forecasts to update inputs across phases and setpoints. Together, these extensions will expand LEMOS beyond batch culture and improve robustness across hosts and circuits.

## Materials and Methods

### LEMOS Device and PDMS Microwell Fabrication

The LEMOS device integrates the electronic components necessary for optical stimulation and wireless communication with an external computer running a Python script. It is built on a 3D-printed PLA frame designed to fit within a standard single-well plate and currently supports 16 microwells with independently controlled LEDs. A full list of components is provided in the SI Section 1. The 16 LEDs in the LEMOS device are connected via a single I^2^C bus controlled by an Arduino Nano33 IoT microcontroller, which also includes a Bluetooth module for wireless communication with an external computer. Both the microcontroller and LEDs are powered by an onboard Li-ion battery. Each LED is individually addressable over the I^2^C bus, with user-defined control over color and intensity. The emitted wavelengths are determined by the number and type of discrete diodes in each multi-color LED. In this design, we selected commercial LEDs containing three diodes with emission peaks at 620–625 nm, 522–525 nm, and 465–467 nm. Compatible with the regulating spectra (535 nm and 650 nm) of the CcaSR system. These LEDs use pulse-width modulation (PWM) for intensity control, which limits analog tuning. As a result, a fixed intensity was chosen to balance sufficient optical stimulation with minimal heat generation.

PDMS microwells were fabricated in the LEMOS device using standard soft lithography techniques. A 1:10 ratio of the curing and base agents of RTV 615 (Momentive Inc.) was mixed thoroughly and degassed under vacuum in a desiccator for up to one hour to remove air bubbles. It is critical to remove the air bubbles from the PDMS before casting since the bubbles change the optical transmission properties of the bottoms of each well leading to non-uniform baseline OD measurement across wells. The degassed PDMS mixture was then poured into the LEMOS device, into which a 3D-printed PLA mold was inserted to define the shape of the wells. The PDMS in the device was cured overnight at room temperature, after which the PLA mold was carefully removed. After each experiment the culture in the PDMS microwells were discarded along with the PDMS to reuse the LEMOS device and fabricate new PDMS microwells.

### Experimental Software Programming

LEMOS operation requires data readout from the microplate reader after each control period (ten minutes). On a computer running Biotek Gen5 (the software interfacing the Biotek microplate reader), a custom Python script monitors and collects the exported data after each cycle, storing fluorescence (FL) and OD_600_ values in separate CSV files. If the CSV files do not exist, the script creates them automatically. To avoid export errors from the microplate reader, the script deletes each exported file after processing. In addition to data handling, the script also ensures the microplate reader remains active by issuing appropriate commands, and it performs the necessary calculations for closed-loop control (see GitHub for implementation details).

### Strains and Culture

The experimental strain was constructed by transforming *E. coli* MG1655 with plasmid pNO286-3 (Addgene #107746) and plasmid pSR58.6 (Addgene #63176). As a negative control for autofluorescence, we constructed an empty backbone by deleting the functional gene cassettes, including P_cpcG_2 and sfGFP, using standard inverse PCR. Cells were plated on LB agar containing 50 μg/mL spectinomycin and 25 μg/mL chloramphenicol and incubated overnight at 37 °C, after which the plates were stored at 4 °C in the dark to minimize light exposure. All experiments were performed in Minimal M9CA Broth (Teknova #M8010) supplemented with 50 μg/mL spectinomycin and 25 μg/mL chloramphenicol. For each condition, a single colony was inoculated into 2 mL of media in a sterile culture tube and grown overnight at 37 °C with shaking at 220 rpm. To prevent unintended light exposure, all cultures were maintained in foil-wrapped tubes. A 100 µL aliquot of the overnight culture was transferred into 2 mL of fresh media in a sterile culture tube and incubated for 4 hours at 37 °C with shaking at 220 rpm. By the end of this period, cultures reached an optical density (OD_600_) of 0.4–0.6. Experimental cultures were then prepared by diluting the pre-culture to an OD_600_ of 0.1 and 200 µL of each condition was added to the PDMS sleeves of the LEMOS device. The device was sealed with a Breath-Easy film (USA Scientific #9123-6100) and loaded into the microplate reader to initiate the experiment. Fluorescence (excitation: 485 nm; emission: 528 nm; gain: 50) and optical density (OD_600_) were measured every 10 minutes using a BioTek Synergy H1 microplate reader, incubated at 30 °C with maximum linear shaking.

### Statistical Analysis

All single colonies were picked randomly from agar plates where the cells were transformed with purified circular DNA using antibiotics as selection markers. No manual group allocation methods were used. Each plate resulted from an independent transformation, and all colonies were assumed to be biological replicates. No data was excluded from the analyses in experiments presented in Fig. 1E, 1F, 2B, 2C, 2E, 3B, 3C, 3D, 4A, 4B, 5B, 5D, 5E, 5F, 5I, 5K, S7 and S8.

## Acknowledgments

The authors would like to thank Richard M. Murray for their insightful discussions. This research is partially supported by the Institute for Collaborative Biotechnologies (ICB) through contract W911NF-19-D-0001 from the U.S. Army Research Office. In addition, the author K.P. was also supported by the Caltech Summer Undergraduate Research Fellowships (SURF) program and the Goldwater Foundation. The authors H.R.N. and B.J. were supported by Texas A&M Engineering Experiment Station.

## Author contributions

K.P., C.Y.H., and A.E. conceptualized the project; K.P. designed and built the LEMOS device and wrote the software for LEMOS-PC interface. H.R.N. and C.Y.H. built the ODE models and designed the controllers; B.J., H.R.N., K.P., and C.Y.H. performed the experiments; H.R.N. and B.J performed data analysis; C.Y.H., H.R.N., B.J., and K.P. wrote the manuscript.

## Data and materials availability

All data needed to evaluate the conclusions in the paper are present in the paper and/or the Supplementary Materials. The source code for all the simulations is available at the following GitHub repository: https://github.com/hariKRN2000/LEMOS-models-and-data. Information about the LEMOS device can be found publicly at the following GitHub repository: https://github.com/kpochana/LEMOS.

