## supplementary Info-v2 for "Closed-loop Optogenetic Control in a Microplate Reader"

Bill of Materials

**Table S1.** Bill of materials for LEMOS device construction. Create a page break and paste in the Table above the caption.

| Sr.no | Item | Link | Notes |
|---|---|---|---|
| 1 | Arduino Nano33 IoT | https://www.amazon.com/Arduino-Nano-33-IoT/dp/B07VW9TSKD | One runs the device, other is used with the central computer |
| 2 | Lipo battery | https://www.amazon.com/gp/product/B0BXNDNTRP | Battery for device |
| 3 | Battery management board | https://www.amazon.com/gp/product/B071RG4YWM | Handles battery charging |
| 4 | Voltage converter board | https://www.amazon.com/gp/product/B09D3G96KZ | Steps battery voltage to logic-level voltage |
| 5 | WS2812B LED strip | https://www.amazon.com/gp/product/B07BTTY4FL | LEDs for optical stimulation |
| 6 | DP3T switch | https://www.amazon.com/uxcell-Horizontal-Switch-Terminals-Latching/dp/B07H3RPD23 | Power and mode switch for device |
| 7 | 22 AWG solid core wire | https://www.amazon.com/TUOFENG-Electronic-Prototyping-Circuits-Breadboarding/dp/B07TX6BX47 | Hookup wire |
| 8 | SUNLU 1.75mm PETG 1kg - Black | https://www.amazon.com/SUNLU-Official-Elite-Filament-1-75mm/dp/B0CFLW4LCJ | Black PETG filament for device frame |

Running LEMOS experiment

**Gen5 operation:**

a) LEMOS experiment initialization. Open the ‘closed loop protocol’ Gen5 file.

b) Select ‘Create experiment and read now’ (Fig. S1) and save the experiment file in the folder where the FL and OD values will be exported (normally the file named Datafile within the experiment directory).

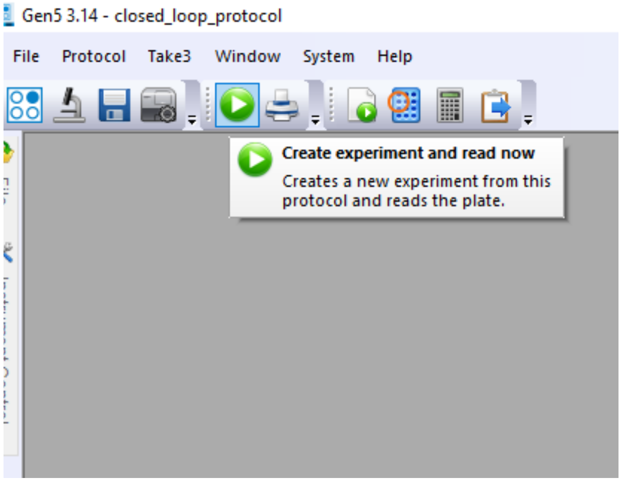


**Fig. S1 Create experiment for LEMOS**

c) Specify the location of the file exported from Gen5 (Fig. S2).

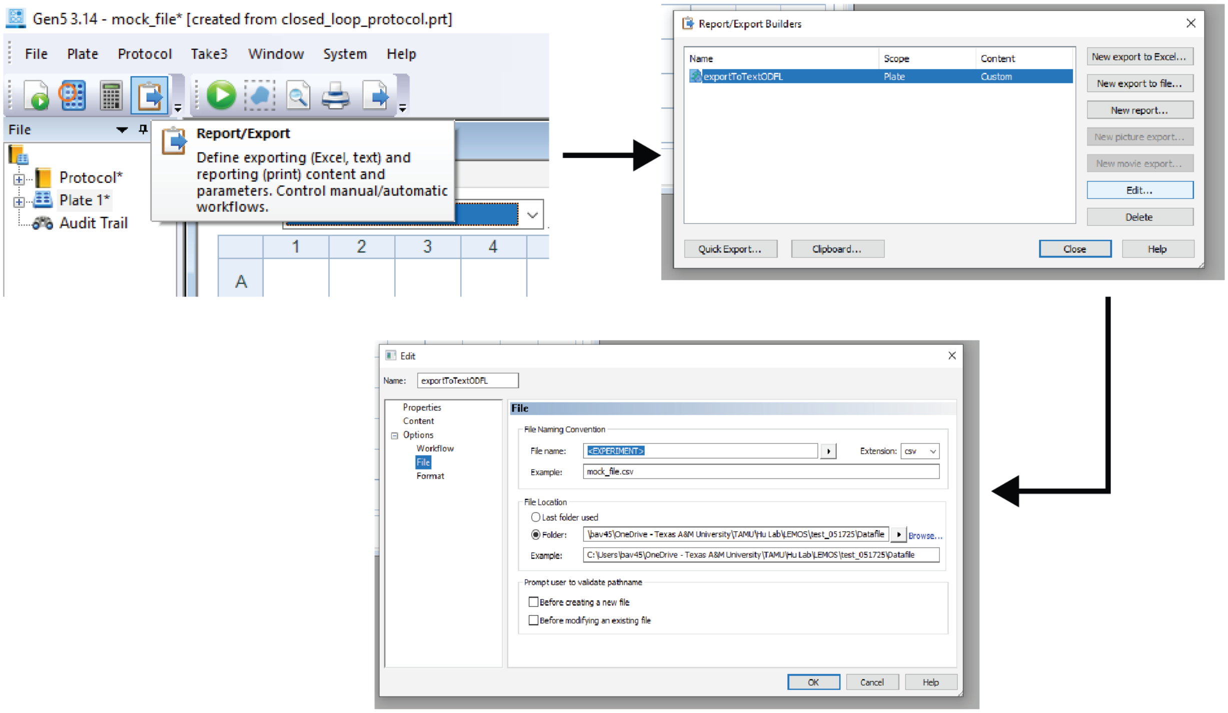


**Fig. S2 Choosing the directory for experiment data export**

d) Click on ‘Procedure’ to change the incubation temperature if required (Fig. S3).

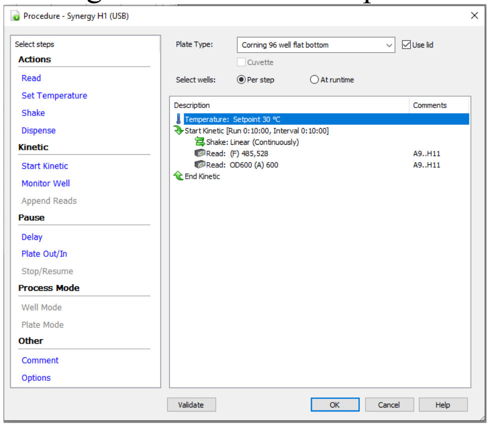


**Fig. S3 Changing the incubation temperature**

e) Click on ‘Run’, wait for the microplate reader to reach the desired incubation temperature, do not hit override or that will produce an error in the python script (Fig. S4).

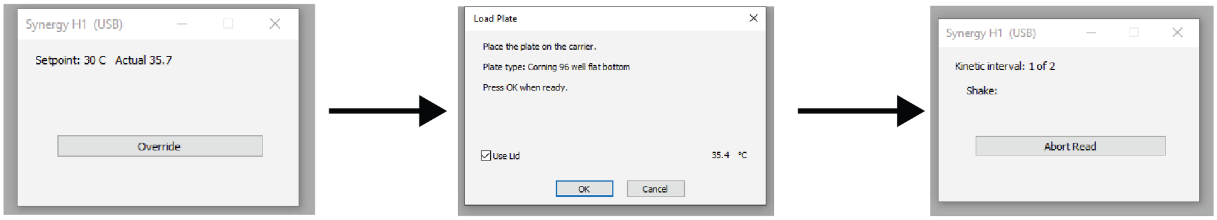


**Fig. S4 Running the LEMOS experiment protocol**

f) Once the microplate reader is at the desired temperature click ‘OK’.

g) Once the study starts running, hit ‘Abort Read’. This step has to be done only once in the beginning for the Gen5 to prompt the ‘Continue Reading Plate 1’ whenever ‘Run’ is selected by the python script (Fig. S5).

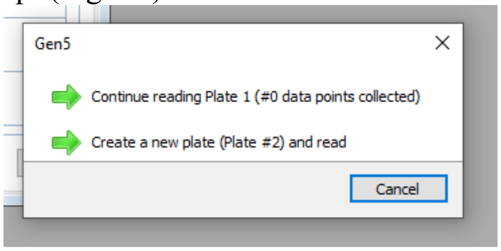


**Fig. S5 Prompt to continue the reading in automated way**

### Python operation:

h) Select the folder where the data from the Gen5 software will be exported. This folder should be updated in the python script accordingly as described earlier (Fig. S6).

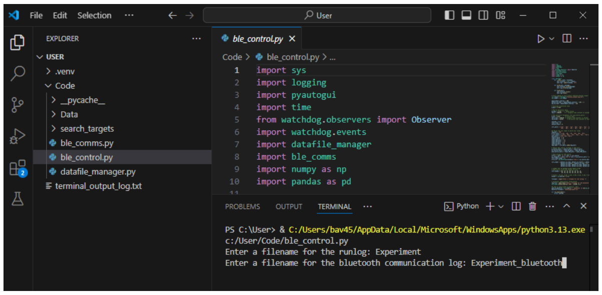


**Fig. S6 Setting up the right directory location in Python script**

i) Launch the python script

Mono-Scale Model

To investigate the consistent overshoot observed under P-control regulation, we developed a deterministic kinetic model to capture the system's dynamic behavior. We used a Chemical Reaction Network (CRN) to represent the signal sensing and transduction dynamics of the two-component optogenetic system, and a set of ordinary differential equations (ODEs) to model the downstream gene expression process.

The kinetics of the sensing component is described with the following CRNs:

$$CCaS \underset{k_{rvrs}}{\overset{k_{fwd}}{\rightleftarrows}} CcaS_P \qquad S1$$

$$CcaS_P + CcaR \rightleftarrows CcaS + CcaR_P \qquad S2$$

$$CcaR_p + CcaR_p \rightleftarrows Ac \qquad S3$$

$$k_{fwd} = k_{green} + k_{fb} \qquad S4$$

$$k_{rvrs} = k_{red} + k_{rb} \qquad S5$$

These reactions are shown in the schematic Fig. 2a. The sensing component of the optogenetic system begins with the transmembrane sensor $CcaS$, which becomes phosphorylated ($CcaS_P$) when exposed to green light. $CcaS_P$ then transfers the phosphate group to the response regulator $CcaR$, producing phosphorylated $CcaR$ ($CcaR_p$), which dimerizes to form the active transcription factor complex $Ac$. This activating complex initiates transcription of the sfGFP gene. To model light-dependent phosphorylation of CcaS, we defined the forward rate constant in equation (S1) as the sum of a small basal phosphorylation rate ($k_{fb}$) and the green-light–dependent rate ($k_{green}$) (equation (S4)). Under green light, phosphorylation occurs at the full rate $k_{green} + k_{fb}$, whereas in its absence (red light or darkness), only the basal rate applies ($k_{fb}$). Similarly, the $CcaR_p$dephosphorylation rate constant was defined as the sum of a basal dephosphorylation rate $k_{rb}$ and the red-light–dependent rate ($k_{red}$), as described in equation (S5). Under red light, dephosphorylation occurs at the full rate $k_{red} + k_{rb}$, while in its absence (green light or darkness), only the basal rate ($k_{rb}$) is applied. The downstream processes of transcription, translation, and protein maturation are captured by the following system of ODEs described in equations S11-S13 (see SI section 3.1 for complete model equations).

Here, $Kc$ is the dissociation constant for the activating complex $Ac$ activating the $P_{cpcG2}$ promoter. The gene expression dynamics were modeled using the standard gene expression modeling framework(*1*), containing three main species including mRNA ($M$), unfolded sfGFP ($P$), and folded sfGFP ($P_m$). In this system, $\beta$ and $k_{tl}$ represent the transcription and translation rates, respectively; $d_m$ denotes the mRNA degradation rate; $d_p$ accounts for the protein dilution; and $k_{fold}$ represents the protein folding rate.

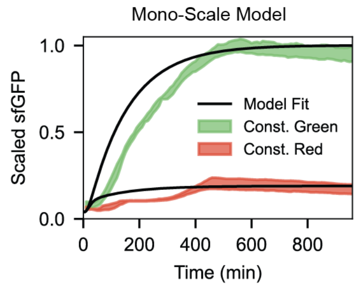


**Fig. S7** Mono-scale model simulation of scaled sfGFP under constant green light activation and constant red-light repression, overlaid with experimental data (N=4, n=2); black solid line indicates the model fit. N denotes the number of technical replicates; n denotes for the number of biological replicates.

Model ODEs

$$\frac{dCcaS}{dt} = -\left(k_{green} + k_{fb}\right) \cdot CcaS + \left(k_{red} + k_{rb}\right) \cdot CcaS_P + k_{R_b} \cdot CcaS_p \cdot CcaR - k_{R_u} \cdot CcaS \cdot CcaR_P \quad S6$$

$$\frac{dCcaS_p}{dt} = \left(k_{green} + k_{fb}\right) \cdot CcaS - \left(k_{red} + k_{rb}\right) \cdot CcaS_P - k_{R_b} \cdot CcaS_p \cdot CcaR + k_{R_u} \cdot CcaS \cdot CcaR_P \quad S7$$

$$\frac{dCcaR}{dt} = -k_{R_b} \cdot CcaS_p \cdot CcaR + k_{R_u} \cdot CcaS \cdot CcaR_P \quad S8$$

$$\frac{dCcaR_p}{dt} = k_{R_b} \cdot CcaS_p \cdot CcaR - k_{R_u} \cdot CcaS \cdot CcaR_P - k_{Rp_b} \cdot CcaR_p^2 + k_{Rp_u} \cdot Ac \quad S9$$

$$\frac{dAc}{dt} = k_{Rp_b} \cdot CcaR_p^2 - k_{Rp_u} \cdot Ac \quad S10$$

$$\frac{dM}{dt} = \beta_m \cdot \left(\frac{Ac}{Kc + Ac} + l_0\right) - d_m \cdot M \quad S11$$

$$\frac{dP}{dt} = k_{tl} \cdot M - \left(d_p + k_{fold}\right) \cdot P \quad S12$$

$$\frac{dP_m}{dt} = k_{fold} \cdot P - d_p \cdot P_m \quad S13$$

Model species

Species explicitly modeled in the process:

**Table S2** Growth independent model species

| Species | Description |
|---|---|
| $M$ | mRNA coding for sfGFP |
| $P$ | Unfolded sfGFP |
| $P_m$ | Folded sfGFP |
| $CcaS$ | CcaS (membrane protein) |
| $CcaS_P$ | Phosphorylated CcaS |
| $CcaR$ | CcaR (response regulator protein) |
| $CcaR_P$ | Phosphorylated CcaR |
| $Ac$ | Transcription activation complex |

Model parameters

**Table S3** Growth independent model parameters

| Parameter | Description | Unit | Value |
|---|---|---|---|
| $\beta_m$ | Transcription rate per plasmid | $nM \cdot min^{-1}$ | 1 |
| $l_0$ | Leak coefficient of promoter | $N/A$ | 1.5e-1 |
| $K_c$ | Dissociation constant of Ac binding to promoter | $nM$ | 4 |
| $d_m$ | mRNA degradation rate constant | $min^{-1}$ | 1e-1 |
| $k_{tl}$ | Translation elongation rate | $min^{-1}$ | 1 |
| $d_p$ | Protein degradation rate | $min^{-1}$ | 7e-3 |
| $k_{fold}$ | sfYFP maturation rate | $min^{-1}$ | 1e-1 |
| $k_{green}$ | Phosphorylation rate of CcaS under green light | $min^{-1}$ | 1 |
| $k_{fb}$ | Basal phosphorylation rate of CcaS | $min^{-1}$ | 1e-1 |
| $k_{red}$ | Dephosphorylation rate of CcaS under red light | $min^{-1}$ | 8e-1 |
| $k_{rb}$ | Basal dephosphorylation rate of CcaS under red light exposure | $min^{-1}$ | 4e-1 |
| $k_{R_b}$ | Phosphorylation rate of CcaR by $CcaS_p$ | $nM \cdot min^{-1}$ | 5e-2 |
| $k_{R_u}$ | Dephosphorylation rate of $CcaR_p$ | $nM \cdot min^{-1}$ | 2.5e1 |
| $k_{R_{pb}}$ | Forward dimerization rate of $CcaR_p$ | $nM \cdot min^{-1}$ | 100 |
| $k_{Rp_u}$ | Reverse dimerization rate of $CcaR_p$ | $min^{-1}$ | 0.5 |

GEAGS model

To incorporate both growth and gene expression dynamics, we applied our previously established dual-scale Gene Expression Across Growth Stages (GEAGS) model framework(*2*). In this model, equations (S18-S21) replace the earlier gene expression equations (S11–S13), while the phosphorylation reactions equations (S1–S3) remain unchanged. The dual-scale model combines ordinary differential equations (ODEs equations (S18-S21)) and chemical reaction network (CRN) equations (S14–S15) to capture gene expression dynamics (see SI section 4.1 for complete model equations), with an additional growth equation (equation (S22)) representing cell population dynamics.

In the growth dynamics equation (S22), $C$ represents the cell count and $C_{max}$ is the carrying capacity, defined by the nutrient availability and volume of the batch culture. The rate modifying functions (RMFs), $\alpha$, $\delta$ and $\gamma$, capture how gene expression dynamics change as cells transition through different stages of growth. These RMFs are defined in terms of the normalized cell density $f = \frac{C}{C_{max}}$, which reflects the proximity of the population to its carrying capacity $C_{max}$. The first RMF, $\alpha = 1 - f$, represents the effective rate of cell division as biomass accumulates in the batch culture. In the model, $\alpha$ modulates the dilution rates of mRNA as well as both folded and unfolded sfGFP. The 2nd RMF, $\delta = \frac{f^n}{1+f^n}$, where $n$ is the Hill coefficient, influences the steepness of the function $\delta$. In the model, $\delta$ modulates the degradation of both folded and unfolded sfGFP, capturing the upregulation of proteolysis as cells enter stationary phase. The 3rd RMF $\gamma = \left(f \cdot (1-f)\right)^m$, reflects the molecular process efficiency across growth phases, peaking during mid-log when growth is most rapid. In the model, $\gamma$ modulates the effective transcription rate, sfGFP folding rate and the coarse-grained translation resource ($R$) availability. Additional parameters in the growth dependent model include $d_{dil}$, the dilution rate due to cell growth, and $k_{tli_b}$ and $k_{tli_u}$, the binding and unbinding rates of the coarse-grained translation initiation complex $C_{tic}$, respectively. The dynamics of $R$ and $C_{tic}$ are captured by the following equations:

$$R_{free} + M \underset{k_{tli_u}}{\overset{k_{tli_b}}{\rightleftarrows}} C_{TIC} \qquad S14$$

$$C_{TIC} \overset{k_{tl}}{\rightarrow} P + M + R_{free} \qquad S15$$

$$R_{total} = R_{max} \cdot \gamma \qquad S16$$

$$R_{free} = R_{total} - C_{TIC} \qquad S17$$

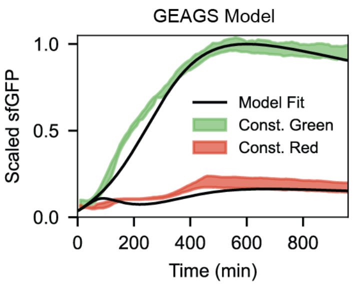


**Fig. S8** Dual-scale GEAGS (Gene Expression Across Growth Stages) model simulation overlaid with the same constant-input data in Fig. S7.

GEAGS Model ODEs

$$\frac{dCcaS}{dt} = -(k_{green} + k_{fb}) \cdot CcaS + (k_{red} + k_{rb}) \cdot CcaS_P + k_{R_b} \cdot CcaS_p \cdot CcaR - k_{R_u} \cdot CcaS \cdot CcaR_P \quad S6$$

$$\frac{dCcaS_p}{dt} = (k_{green} + k_{fb}) \cdot CcaS - (k_{red} + k_{rb}) \cdot CcaS_P - k_{R_b} \cdot CcaS_p \cdot CcaR + k_{R_u} \cdot CcaS \cdot CcaR_P \quad S7$$

$$\frac{dCcaR}{dt} = -k_{R_b} \cdot CcaS_p \cdot CcaR + k_{R_u} \cdot CcaS \cdot CcaR_P \quad S8$$

$$\frac{dCcaR_p}{dt} = k_{R_b} \cdot CcaS_p \cdot CcaR - k_{R_u} \cdot CcaS \cdot CcaR_P - k_{Rp_b} \cdot CcaR_p^2 + k_{Rp_u} \cdot Ac \quad S9$$

$$\frac{dAc}{dt} = k_{Rp_b} \cdot CcaR_p^2 - k_{Rp_u} \cdot Ac \quad S10$$

$$\frac{dM}{dt} = \beta_m \cdot \left(\frac{Ac}{Kc + Ac} + l_0\right) - (d_m + d_{dil}) \cdot M - k_{tli_b} \cdot R_{free} \cdot M + k_{tli_u} \cdot C_{tic} + k_{tl} \cdot C_{tic} \quad S18$$

$$\frac{dC_{tic}}{dt} = k_{tli_b} \cdot R_{free} \cdot M - k_{tli_u} \cdot C_{tic} - k_{tl} \cdot C_{tic} \quad S19$$

$$\frac{dP}{dt} = k_{tl} \cdot C_{tic} - (d_p + d_{dil}) \cdot P - k_{fold} \cdot P \quad S20$$

$$\frac{dP_m}{dt} = k_{fold} \cdot P - (d_p + d_{dil}) \cdot P_m \quad S21$$

$$\frac{dC}{dt} = k_{gr} \cdot C \cdot \left(1 - \frac{C}{C_{max}}\right) \quad S22$$

Growth dependent rates:

Applying RMFs to growth-dependent rates:

$$\boldsymbol{\beta_m} = \beta_m \cdot \gamma \quad S23$$

$$\boldsymbol{k_{fold}} = k_{fold} \cdot (\gamma + b_{fold}) \quad S24$$

$$\boldsymbol{d_m} = d_m \cdot \alpha \quad S25$$

$$\boldsymbol{d_p} = d_p \cdot \delta \quad S26$$

$$\boldsymbol{d_{dil}} = d_{dil} \cdot \alpha \quad S27$$

Conservation equations for growth dependent translation resource (R):

$$R_{total} = R_{max} \cdot \gamma \quad \text{S28}$$
$$R_{free} = R_{total} - C_{tic} \quad S29$$

Model species

**Table S4** GEAGS model species

| Species | Description |
|---|---|
| $M$ | mRNA coding for sfGFP |
| $P$ | Unfolded sfGFP |
| $P_m$ | Folded sfGFP |
| $CcaS$ | CcaS (membrane protein) |
| $CcaS_P$ | Phosphorylated CcaS |
| $CcaR$ | CcaR (response regulator protein) |
| $CcaR_P$ | Phosphorylated CcaR |
| $Ac$ | Transcription activation complex |
| $C_{tic}$ | Translation initiation complex |
| $R$ | Coarse-grained translation resource |
| $C$ | Cell population |

Model parameters

**Table S5** GEAGS model parameters

| Parameter | Description | Unit | Estimate |
|---|---|---|---|
| $\beta_m$ | Transcription rate per plasmid | $nM \cdot min^{-1}$ | 2.8e1 |
| $l_0$ | Leak coefficient of promoter | $N/A$ | 1e-5 |
| $K_c$ | Dissociation constant of Ac binding to promoter | $nM$ | 4.5e1 |
| $d_m$ | mRNA degradation rate constant | $min^{-1}$ | 2.7e-1 |
| $k_{tli_b}$ | $C_{tic}$ formation rate | $nM^{-1} \cdot min^{-1}$ | 4e1 |
| $k_{tli_u}$ | $C_{tic}$ dissociation rate | $min^{-1}$ | 1e1 |
| $k_{tl}$ | Translation elongation rate | $min^{-1}$ | 2 |
| $d_p$ | Protein degradation rate | $min^{-1}$ | 8e-4 |
| $k_{fold}$ | sfYFP maturation rate | $min^{-1}$ | 3e-1 |
| $b_{fold}$ | Basal coefficient for $k_{fold}$ | $N/A$ | 1 |
| $k_{green}$ | Phosphorylation rate of CcaS under green light | $min^{-1}$ | 8e-1 |
| $k_{fb}$ | Basal phosphorylation rate of CcaS | $min^{-1}$ | 3e-2 |
| $k_{red}$ | Dephosphorylation rate of CcaS under red light | $min^{-1}$ | 1.3 |
| $k_{rb}$ | Basal dephosphorylation rate of CcaS under red light exposure | $min^{-1}$ | 8e-1 |
| $k_{R_b}$ | Phosphorylation rate of CcaR by $CcaS_p$ | $nM \cdot min^{-1}$ | 5e-2 |
| $k_{R_u}$ | Dephosphorylation rate of $CcaR_p$ | $nM \cdot min^{-1}$ | 2.5e1 |

| | | | |
|---|---|---|---|
| $k_{R_{pb}}$ | Forward dimerization rate of $CcaR_p$ | $nM \cdot min^{-1}$ | 5.5e1 |
| $k_{Rp_u}$ | Reverse dimerization rate of $CcaR_p$ | $min^{-1}$ | 8e-2 |
| $R_{max}$ | Max. total R availability | $nM$ | 4 |
| $n$ | Exponent of $\gamma$ | $N/A$ | 8.9e-1 |
| $n_{delta}$ | Hill coefficient of $\delta$ | $N/A$ | 5.5 |
| $C_0$ | Initial condition for cell population | $counts$ | 4.69e7 |
| $C_{max}$ | Max. cell population (holding capacity) | $counts$ | 7.14e8 |
| $k_{gr}$ | Logistic growth rate | $min^{-1}$ | 1.1e-2 |

Minimal model description

Model ODEs

$$\frac{dM}{dt} = \boldsymbol{\beta_m} \cdot \left(\frac{Ac}{Kc + Ac} + l_0\right) - (\boldsymbol{d_m} + \boldsymbol{d_{dil}}) \cdot M \qquad S30$$

$$\frac{dP}{dt} = \boldsymbol{k_{tl}} \cdot M - (\boldsymbol{d_p} + \boldsymbol{d_{dil}}) \cdot P \qquad S31$$

$$\frac{dC}{dt} = k_{gr} \cdot C \cdot \left(1 - \frac{C}{C_{max}}\right) \qquad S32$$

Modified rate equations:

$$\boldsymbol{\beta_m} = \beta_m \cdot \gamma \qquad S33$$

$$\boldsymbol{k_{tl}} = k_{tl} \cdot \gamma \qquad S34$$

$$\boldsymbol{d_m} = d_m \cdot \alpha \qquad S35$$

$$\boldsymbol{d_p} = d_p \cdot \delta \qquad S36$$

$$\boldsymbol{d_{dil}} = d_{dil} \cdot \alpha \qquad S37$$

Model species

**Table S6** Minimal model species

| Species | Description |
|---|---|
| $M$ | mRNA coding for sfGFP |
| $P$ | sfGFP |
| $C$ | Cell population |

Model parameters

**Table S7** Minimal model parameters

| Parameter | Description | Unit | Estimate |
|---|---|---|---|
| $\beta_m$ | Transcription rate per plasmid | $nM \cdot min^{-1}$ | 3 |
| $l_0$ | Leak coefficient of promoter | $N/A$ | 5e-3 |
| $K_c$ | Dissociation constant of Ac binding to promoter | $nM$ | 10 |
| $d_m$ | mRNA degradation rate constant | $min^{-1}$ | 2e-1 |
| $k_{tl}$ | Translation elongation rate | $min^{-1}$ | 1.1 |
| $d_p$ | Protein degradation rate | $min^{-1}$ | 1e-4 |
| $n$ | Exponent of $\gamma$ | $N/A$ | 0.9 |
| $n_{delta}$ | Hill coefficient of $\delta$ | $N/A$ | 5.5 |
| $C_0$ | Initial condition for cell population | $counts$ | 4.69e7 |
| $C_{max}$ | Max. cell population (holding capacity) | $counts$ | 8.37e8 |
| $k_{gr}$ | Logistic growth rate | $min^{-1}$ | 1.2e-2 |

Assume steady state values of $Ac$ for each light input:

Under green light: $Ac = 80\ nM$
Under red light: $Ac = 1\ nM$
Under no light (dark): $Ac = 3\ nM$
